# MdAGG apple lectins in Fire Blight resistance: CRISPR/Cas9 validation and their potential for intragenesis approaches

**DOI:** 10.1101/2024.11.08.622591

**Authors:** Antoine Bodelot, Nicolas Dousset, Elisa Ravon, Christelle Heintz, Marie-Noelle Brisset, Alexandre Degrave, Emilie Vergne

## Abstract

Fire blight, caused by the bacterium *Erwinia amylovora*, represents a significant threat to apple (*Malus domestica*) production. Currently, only a limited number of genes effectively involved in resistance to *E. amylovora* have been identified.

Seeking new resistance candidates, we focused on a multigene family encoding amaranthin-like lectins, which are highly upregulated following chemical elicitation by acibenzolar-S-methyl (ASM). These lectins are believed to contribute to downstream defense by promoting bacterial aggregation, which led to their designation as *M. domestica* agglutinins (MdAGG). When loss-of-function editions were introduced into *MdAGG* genes, the plant’s ability to mount a fully effective defense response against fire blight upon ASM treatment was compromised, confirming the role of MdAGGs in fire blight resistance. Next, we coupled the *pPPO16* promoter, endogenous to apple and known to be rapidly induced during *E. amylovora* infection, with the coding sequence of MdAGG10 to create apple lines with fire blight-inducible *MdAGG10* expression. Early *MdAGG10* expression in these lines significantly improved resistance to fire blight, and an additional ASM treatment further enhanced this resistance.

In summary, we conclude that MdAGGs act as defense genes whose timely expression can provide effective resistance against *E. amylovora*.

## Introduction

Fire blight is a devastating disease caused by the necrogenic bacteria *Erwinia amylovora* (*Ea*) which affects most members of the *Rosaceae* family, including apple (*Malus domestica*). *Ea* owns two major virulence factors. The first one is its type III secretion system (T3SS) which injects effectors into host cells and five effectors are assumed to be secreted and translocated via this T3SS *ie.* AvrRpt2_Ea_, DspE/A, HopPtoC_Ea_, Eop1 and Eop3 (McNally *et al*., 2015). The second factor is the exopolysaccharide (EPS), external glycans whose production is correlated to *Ea* virulence (Ayers *et al*., 1979). As a Gram negative bacteria, the external membrane of *Ea* cells also contains lipopolysaccharides (LPS), which are glycolipids protecting the bacterial cells from the oxidative burst launched by the plant during the infection process (Berry *et al*., 2009). EPS and LPS are known bacterial PAMPs (Pathogen-Associated Molecular Patterns), widely conserved components essential for pathogenicity and pathogens survival (Boutrot & Zipfel, 2017).

As the protection practices against fire blight are scarce and often poorly effective (Bonn & Van der Zwet, 2000), the selection of resistant cultivars remains an alternative to be favoured. However, conventional breeding of apple is tedious especially due to a long juvenile phase (Campa *et al*., 2023). Until now, conventional breeding for fire blight resistance has mostly focused on resistance QTLs (Quantitative Trait Loci), with the strongest ones identified in wild *Malus*: on linkage group (LG) 3 of *Malus × robusta 5* (Peil *et al*., 2019), on LG10 of *M. fusca* (Emeriewen *et al*., 2014) and on LG12 of ‘Evereste’, *M. floribunda 821* (Durel *et al*., 2009) and *M. × arnoldiana* (Emeriewen *et al*., 2017). Genes under these QTLs turn out to represent Resistance (R) genes involved in gene-for-gene recognition and subsequent triggering of ETI (Effector Triggered Immunity; Jones & Dangl, 2006). They have been assigned as either NBS-LRR candidate genes: *FB_MR5* in *Malus × robusta 5* (Fahrentrapp *et al*., 2013); receptor-like kinase candidate genes: *FB_Mfu10* in *M. fusca* (Emeriewen *et al*., 2018); both of them being candidates in the QTL on LG12 of ‘Evereste’ (and *M. floribunda*; Parravicini *et al*., 2011) and *M. × arnoldiana* (Emeriewen *et al*., 2021). However, bypasses of the resistance gained from the introgression of QTLs from ‘Evereste’ (and *M. floribunda*) and *M. robusta* (*FB_MR5* gene) into commercial varieties have already been observed in apple, due to the evolution of the bacterial effectors Eop1 and AvrRPT2_Ea_, respectively (Emeriewen *et al*., 2019).

New breeding technologies, including cisgenesis, intragenesis, RNA interference and genome editing mediated by CRISPR-Cas9 are alternatives to conventional breeding (Campa *et al*., 2023). These technologies, however, have rarely been used to improve resistance to fire blight, via R recognition mechanisms or via targeting mechanisms considered to be more durable. Cisgenesis has been used to introduce the *FB_MR5* gene from *Malus × robusta 5* in the susceptible genotype ‘Gala Galaxy’, leading to resistance levels equivalent to those obtained for classically bred descendants. Intragenesis has already been employed to improve scab resistance in apple (Joshi *et al*., 2011), but not yet for fire blight resistance. Successful knockdown of apple susceptibility genes (S-genes; Campa *et al*., 2019; Pompili *et al*., 2020) by has been performed by RNA interference on the *HIPM* gene (Campa *et al*., 2019) or by directed mutagenesis by CRISPR/Cas9 of the *DIPM4* gene (Pompili *et al*., 2020), both genes encoding proteins that respectively interact with *Ea* effectors HrpN and DspA/E. To the best of our knowledge, no published attempts have been made to improve apple resistance to *Ea* through NBT by targeting defense genes, such as Pathogenesis Related (PR) genes, that are involved in downstream mechanisms of pathogen recognition by R proteins or Pattern Recognition Receptors (PRR) and subsequent signal transduction (Pattern-Triggered Immunity, PTI; Jones & Dangl, 2006).

The candidate family we targeted in the present study was identified in preliminary works on ASM-induced resistance in apple against *E. amylovora* and proposed as a new class of PR proteins (Warneys *et al*., 2018; Chavonet *et al*., 2022). ASM up-regulates thousands of genes in apple, including a multigenic family partly organized as tandem array genes encoding lectins that have been termed *Malus domestica* agglutinins (MdAGGs, Warneys *et al*., 2018). Lectins are proteins with at least one non-catalytic site able to reversibly bind mono- or oligosaccharides (Peumans & Van Damme, 1995). In plants, PRR harbouring extracellular lectin domains that recognize specific carbohydrates of pathogens and trigger defense responses have received extensive interest (Bellande *et al*., 2017). In contrast, defense-inducible lectins that are not related to pathogen recognition and subsequent signalization are less studied, although they can bind carbohydrate patterns and act as key players in plant defense processes through different pathways (De Coninck & Van Damme 2021). According to the classification of Van Damme *et al*. (2008) MdAGG1 to MdAGG18 are defense-inducible merolectins composed of a unique amaranthin-like carbohydrate binding domain. The apple genome also encodes nine other amaranthin-like lectins, MdDiAGG1 to 9, composed of two amaranthin-like domains and an additional aerolisin/ETX pore-forming domain for three of them: MdDiAGG5, 6 and 7 (Warneys *et al*., 2018). *MdDiAGGs* are constitutively expressed, unlike *MdAGGs*, and *MdAGG10*, which is one of the most ASM-induced genes and shares the highest percentage of identity with the *MdAGGs* consensus sequence (Warneys *et al*., 2018), has been studied further. Agglutination of *Ea* cells by MdAGG10 has been demonstrated *in vitro*, involving electrostatic interactions with the negatively charged bacterial polysaccharides EPS and LPS. Moreover, *Ea* actively downregulates the expression of *MdAGG* genes and evades MdAGG10-driven cell aggregation thanks to the secretion of EPS (Chavonet *et al*., 2022). *MdAGGs* are therefore assumed to be good candidate defense genes for improving apple PTI against *Ea*. However, there is still no functional evidence supporting the involvement of MdAGGs in apple resistance to *Ea*, as the constitutive overexpression of *MdAGG10* in the susceptible ‘Gala’ genotype did not improve resistance to *Ea* (Bodelot *et al*., 2023).

In this study, thanks to CRISPR/Cas9, now widely mastered in apple (Campa *et al*., 2023) we were able to demonstrate that *MdAGGs* are involved in apple resistance against *Ea*. In addition, using the intragenic association between the fire blight inducible *pPPO16* promoter (Gaucher *et al*., 2022) and the coding sequence of *MdAGG10*, we have further created a partial resistance to fire blight.

## Results

### Generation of transgenic lines

To examine *MdAGG* genes role in apple resistance toward *Ea*, targeted knockouts were generated in the susceptible variety ‘Gala’ using CRISPR/Cas9 editing. To maximize the probability to target the 18 *MdAGG* genes we used a T-DNA construct harboring two guide-RNAs that were designed so that either one or both are complementary to each *MdAGG* gene, while minimizing off-target sequences (Table 1, only *MdDiAGG4* is a potential off-target of gRNA1). We speculated that this strategy would lead to (i) punctual mutations at either gRNA site, (ii) deletions of several hundreds of nucleotides between both gRNA sites within *MdAGG* genes or (iii) deletions of several thousands of nucleotides between gRNA sites of different *MdAGG* genes organized in tandem array.

**Table 1.**
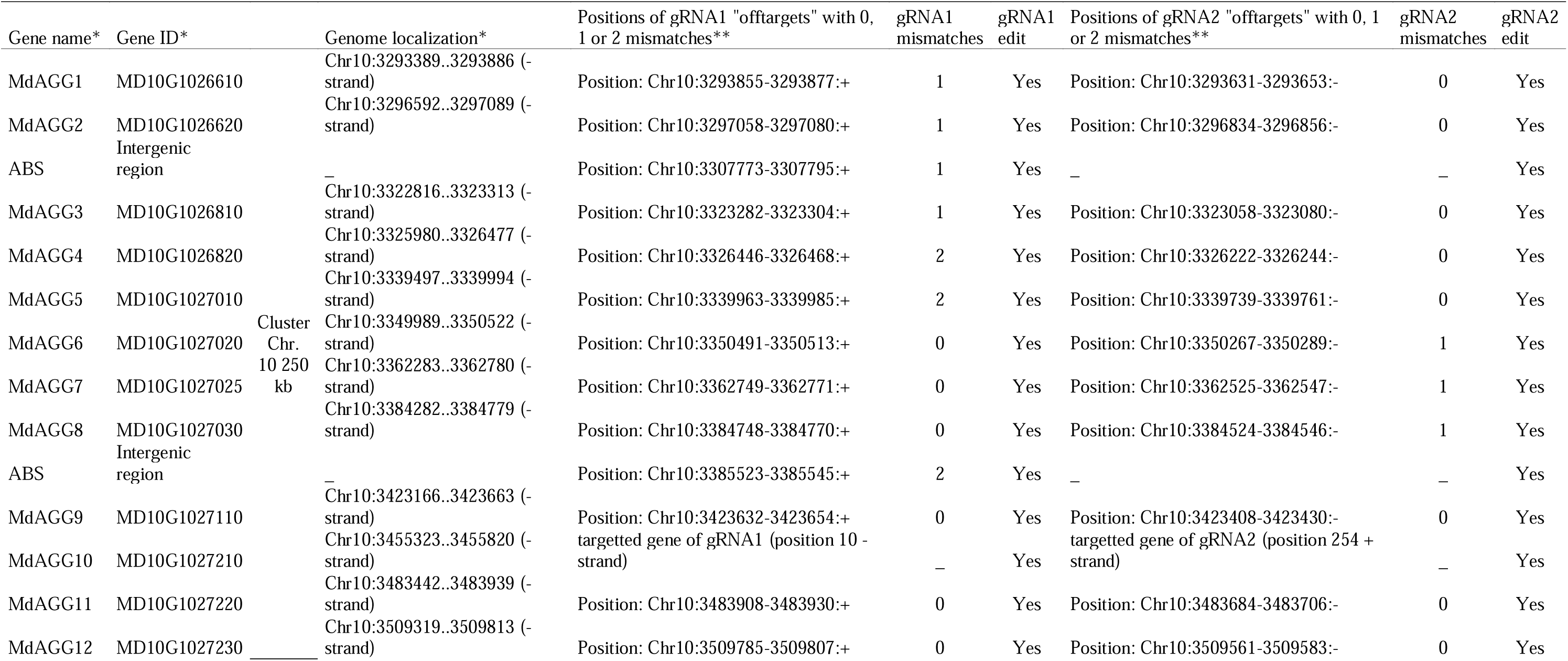
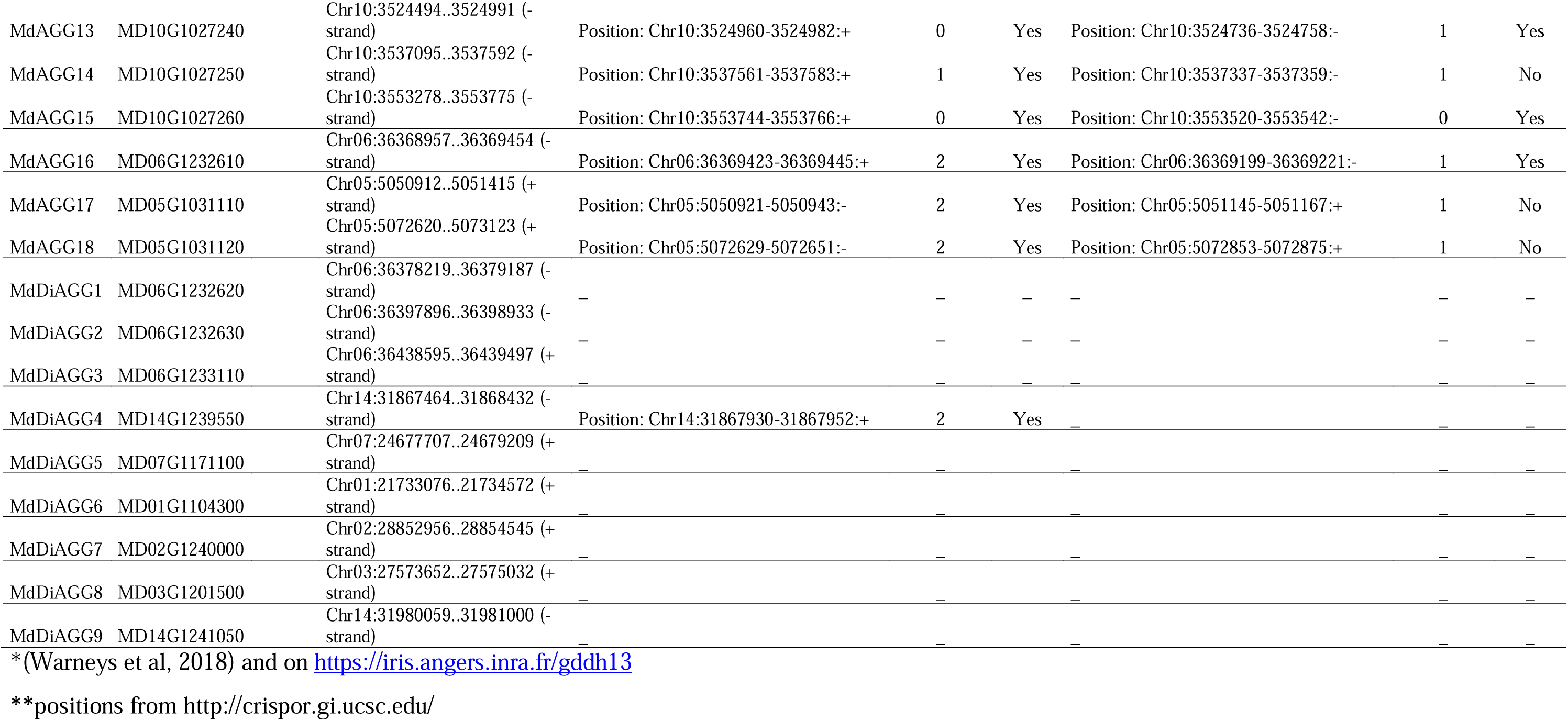
Localization summary of the gRNA1 and gRNA2 targeted DNA zones. gRNAs target all *MdAGG* members of the family. gRNA1 and 2 were designed on *MdAGG10* (MD10G1027210). gRNA1 targets all others *MdAGGs* with more or less mismatches, but also the off-target *MdDiAGG4* (MD14G1239550) with two mismatches. gRNA2 targets all others *MdAGGs* with more or less mismatches except MdAGG14 (MD10G1027250), MdAGG17 (MD05G1031110) and MdAGG18 (MD05G1031120); it has no predicted off-target. Mismatch tolerance is considered effective (“Yes”) for up to two base mismatches (Charrier et al, 2019), occurring between positions 20 and 4 relative to the PAM.

A total of 419 leaf explants were inoculated with *Agrobacterium tumefaciens* containing the binary vector with the CRISPR-MdAGGAll construct (T-DNA in Fig. S1a) targeting supposedly all of *MdAGG* family members (Table 1). The five kanamycin resistant-regenerants which survived over 6 months of micropropagation were analyzed by PCR to confirm transformation and absence of *A. tumefaciens* contamination (Fig. S2a). Amplification using primers I (Table S1) designed for *EF1*α gene was used as a positive PCR control, plasmid DNA extracted from *A. tumefaciens* strain containing the CRISPR-MdAGGAll construct as a positive control of T-DNA presence and genomic DNA extracted from a non-transgenic ‘Gala’ as a negative control of T-DNA-presence. K primers (Table S1) amplifying 23S ribosomal RNA from *A. tumefaciens* showed that all the tested lines were free from *A. tumefaciens* contamination. Amplification using J primers (Table S1), which target the *nptII* selection gene, confirmed that all five lines were true transgenics containing the selectable marker. Finally, amplification using primers B and L (Table S1), which respectively target gRNAs sequences and the Cas9 coding sequence, demonstrated that all lines had integrated the full construct. Considering the initial number of explants (419), the transformation rate was therefore 1.19%. The five lines were called *mdagg-1* to *mdagg-5* and further analyzed.

A previous study showed that constitutive overexpression of *MdAGG10* in the susceptible ‘Gala’ variety does not lead to enhanced resistance toward *Ea* (Bodelot *et al*., 2023). We wondered whether plants engineered to conditionally express *MdAGG10* following *Ea* infection could display partial resistance toward fire blight. The promoter of the endogenous apple *PPO16* gene, known to be activated by *Ea* infection in a rapid and sustained manner (Gaucher *et al*., 2022), was therefore used to produce transgenic *pPPO16::MdAGG10* apple lines. A total of 1150 leaf explants were inoculated with *A. tumefaciens* containing the binary vector with the *pPPO16::MdAGG10* construct (T-DNA in Fig. S1b) assumed to induce MdAGG10 expression upon *Ea* infection (Gaucher *et al*., 2022). The nine kanamycin resistant-regenerants which survived over 6 months of micropropagation were analyzed by PCR, with similar controls as describe above for CRISPR lines, to confirm transformation and absence of *A. tumefaciens* contamination (Fig. S2b). Amplification with primers H (Table S1) for the *pPPO16::MdAGG10* association presence showed that all lines had integrated the full construct. Considering the initial number of explants (1150), the transformation rate was therefore 0.78%. Six of the nine lines, called *pPPO16::MdAGG10-1* to *pPPO16::MdAGG10-6* were further analyzed.

### Dual guide-RNA edition of *MdAGG* genes leads to multiple loss-of-function mutations

For each of the five lines, named *mdagg-1* to *mdagg-5*, four bacterial clones (referred to as ‘clones’) were sequenced. Each clone contained one allele of the potentially edited *MdAGG* target sequence, including the two gRNA target sites. These sequences were obtained through cloned bulk PCR using primers M (Table S1), which flanked the target sequence in all 18 *MdAGG* genes (Fig. 1). Thanks to this partial but qualitative characterization, the two target sites proved effective, and the five lines proved edited, displaying various types of edits, as previously hypothesized: (i) indel or substitution of several nucleotides in one *MdAGG* sequence (called after type A edit), (ii) deletion of the sequence between the two target sites, of around two hundred nucleotides, in one *MdAGG* sequence (called after type B edit) and (iii) deletion of several thousands of nucleotides between two different *MdAGGs* sequences (called after type C edit). This characterization also reported the insertion of one *MdAGG* sequence between the two targeted sites of another *MdAGG* sequence (called after Type D edit). In *mdagg-1* line, we observed four type A edits. In *mdagg-2* line, we observed two type B edits (clone 1 and 2), one type A edit (clone 3) and one type C edit (clone 4) displaying a 126 kb deletion. In *mdagg-3* line, we observed three unedited *MdAGG* sequences (clones 2, 3 and 4) and one type A edit (clone 1). For *mdagg-4* line, clones 1 and 2 displayed type A edits, clone 3 displayed a type C edit with a 212 kb deletion. Ultimately, clone 4 displayed a type D edit, where 200 nucleotides of *MdAGG15* were invertedly inserted between the two targeted sites of *MdAGG3*. For *mdagg-5* line, we observed three type B edits (clones 1, 2 and 3), as well as one type A edit (clone 4). All these edits on *MdAGGs* coding sequence led to the early insertion of a stop codon resulting in a truncated protein, except for one *MdAGG10* allele (clone 3) of *mdagg-1* line, where edits led to substitution or indels of amino acids (Fig. 1). Altogether, this analysis showed that both gRNAs were effective and that they led to multiple loss-of-function mutations in the *MdAGG* genes.

**Fig. 1.**
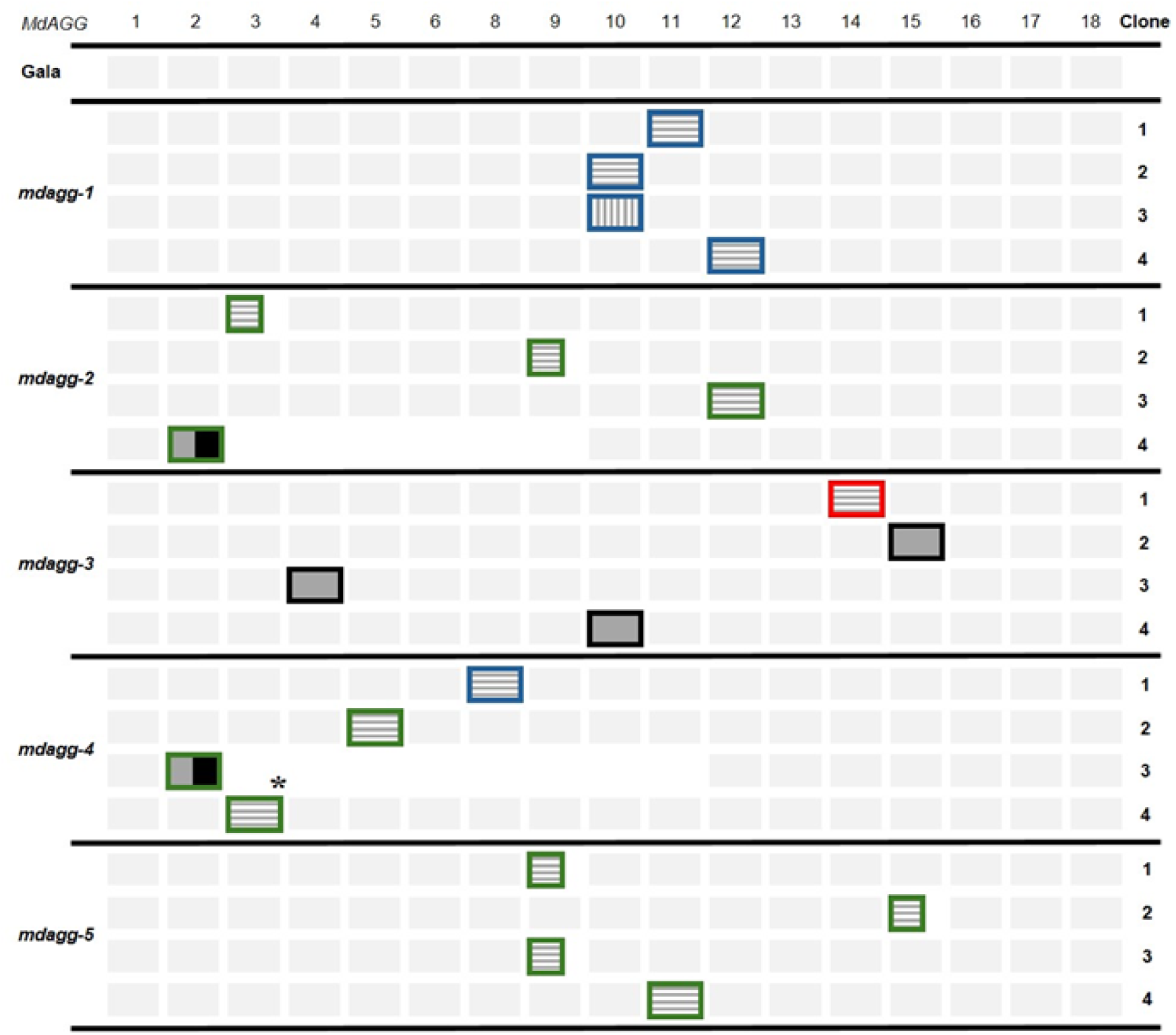
Dual guide-RNA gene edition strategy leads to *MdAGG* genes knockdown in apple. Four cloned sequences from each of the 5 independant *mdagg* edited apple mutants were analyzed. Each box represents one *MdAGG* sequence. Unedited *MdAGG* sequences are represented by a darkgrey box, short indels leading to frameshift mutations protein sequence (respectively early insertion of stop codon) are represented by darkgrey boxes with vertical lines (respectively with horizontal lines), larger deletions occuring within a single *MdAGG* sequence and leading to frameshift mutations are represented with a smaller darkgrey box with grey horizontal lines, deletions occuring between different *MdAGG* sequences and leading to large genomic DNA excisions are represented with a darkgrey and black box, sequences for which no information is available are represented by lightgrey boxes. « * » corresponds to a specific case observed in clone 4 of *mdagg-4* where a 200 nucleotides portion of *MdAGG15* was invertedly inserted into the targeted site of *MdAGG3*. gRNAs efficencies are represented in (i) blue if the edits occured in the gRNA1 targeted sequence, in (ii) red if edits occured in gRNA2 targeted sequence and in (iii) green if edits occured in both gRNAs targeted sequences.

### MdAGG loss-of-function mutants are impaired in their ASM-induced resistance response toward *E. amylovora*

To assess the effect of loss-of-function mutations of *MdAGGs* in inducible-resistance, *in vitro* shoot cultures of lines *mdagg-1* to *5* were treated with ASM and further inoculated with *Ea*. The untransformed susceptible variety ‘Gala’ (WT), either treated with water or ASM, was included as a respectively susceptible and resistant control. Fire blight disease symptoms were scored daily over 8 days following inoculation and resistance was evaluated by determining disease incidence at the end of the experiment or disease severity by AUDPC calculation. As expected, water-treated WT displayed significantly higher disease incidence and severity scores than ASM-treated WT. Compared to the resistant control, fire blight incidence was significantly higher in all *mdagg* mutant lines treated with ASM, except for line *mdagg-4* (Fig. 2). Regarding fire blight severity, the AUDPC of ASM-treated *mdagg-4* mutants was equivalent to that of the resistant control whereas the AUDPC of the four other ASM-treated *mdagg* mutants was equivalent to that of the susceptible control (Fig. 2). Altogether, these results show that except for line *mdagg-4*, *in vitro* shoots of *mdagg* mutant lines were not able to mount a fully-effective resistance response toward *Ea* upon ASM treatment. Mutant lines *mdagg-1* and *mdagg-2* were further analyzed.

**Fig. 2.**
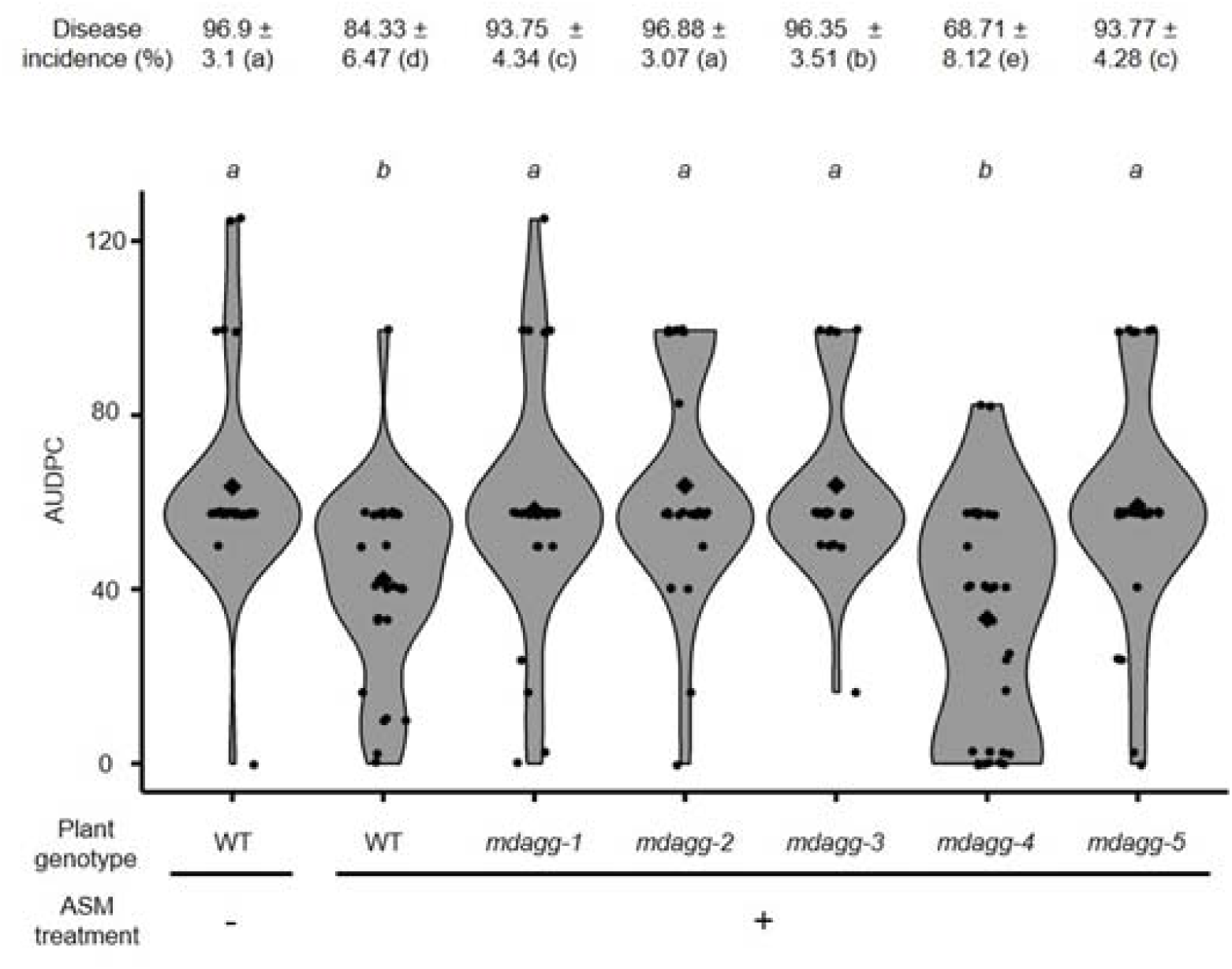
*In vitro* culture shoots of *MdAGGs* edited apple mutants except *mdagg-4* are still susceptible to *Erwinia amylovora* despite ASM treatment. Fire blight disease incidence (± standard deviation) and AUDPC scoring over a 8 day period after *Ea* infection. Each point represents the AUDPC value for individual shoots (28<n< 32), diamonds represent the mean AUDPC for each condition. Significance was evaluated with a Kruskall-Wallis test followed by a Dunn test, different letters indicate different statistical classes (P<0.05).

We completed the quantitative characterization of the edits in *mdagg-1* and *mdagg-2* using a qPCR-based method we developed to measure the editing rates at the gRNA1 targets consisting of MdAGGs target sites and the MdDiAGG4 off-target site (Table 1). The forward primers for both the MdAGGs targets and MdDiAGG4 off-target were designed to end just after the Cas9 cut site (4 nucleotides from the PAM) of the gRNA1 target. This design allows amplification only in wild-type alleles, where no cutting or modification has occurred, and in rare cases where synonymous repairs may have taken place. Estimated global editing rates were then calculated as inverses of these quantities of unmodified sequences. The results show that both mutant lines *mdagg-1 and -2* were significantly edited at the *MdAGG*s target sites, while neither displayed significant edits at the off-target site *MdDiAGG4*. (Fig. 3a).

**Fig. 3.**
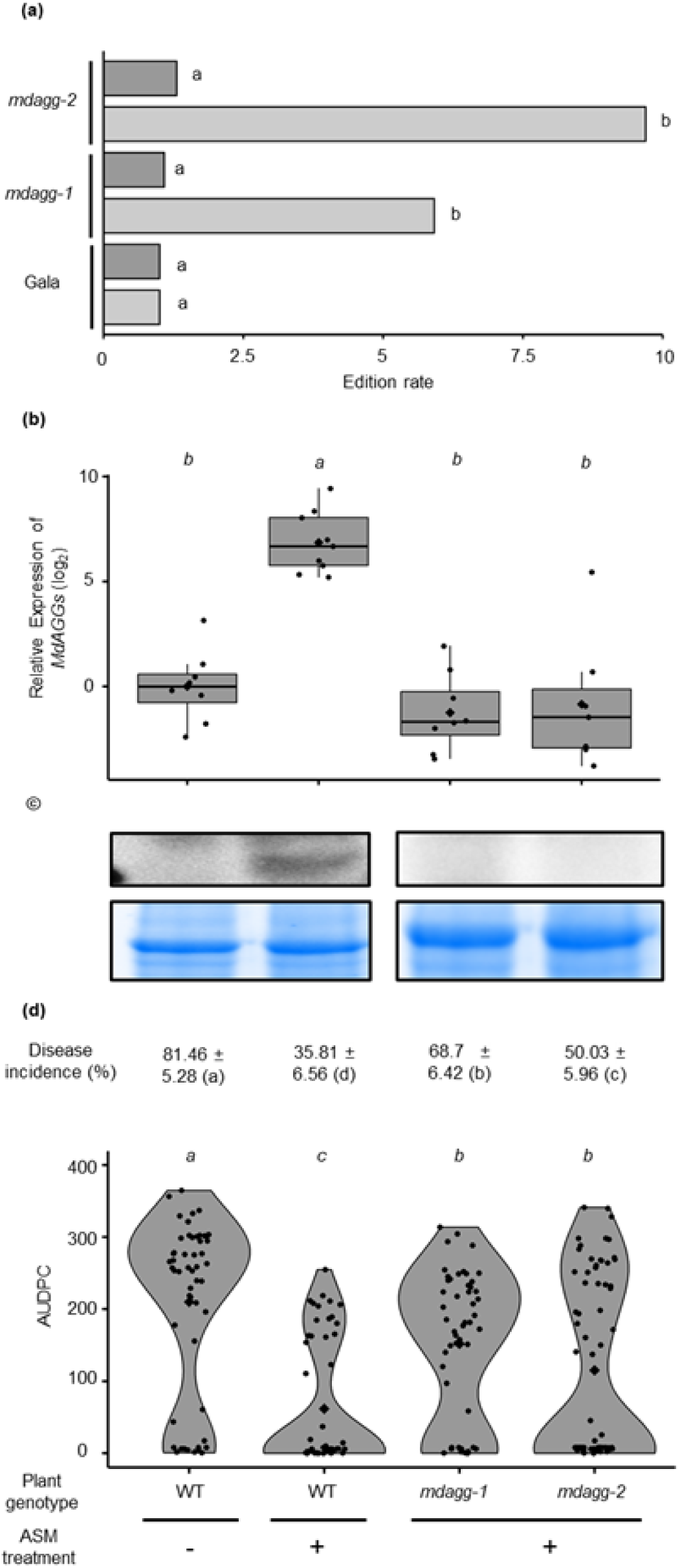
Greenhouse-grown *MdAGG* edited apple mutants do not express *MdAGG* upon ASM treatment and are more susceptible to *Erwinia amylovora*. **(a)** Quantification of editing rates at the gRNA1 target sequences in *mdagg-1* and -*2* lines. The editing rate (e) was calculated as followed: e = 1/x where x corresponds to the relative quantity of unedited DNA sequences of *MdAGGs* or *MdDiAGG4* measured by qPCR. The values shown are the average measured in 4 to 6 samples from two independant biological repeats. The data are compared with the ‘Gala’ non transformed genotype, as an unedited reference. Significance of the Wilcoxon rank test: *P<0.05; ns not significant. **(b)** Relative expression (log_2_ ratio) of *MdAGGs* in the inoculated leaves at the time of bacterial infection. Means, medians, and interquartile range (IQR) are indicated by diamonds, lines, boxes, respectively. Whiskers indicate the most extreme values between the nearest quartile and 1.5 IQR. The mean value of control-treated untransformed (WT) plant samples was used for the calculation of the log_2_ ratio. Each dot represents one biological replicate*, i.e* a pool of two leaves of independant plants (7<n<9). **(c)** MdAGGs protein detection by Western Blot (upper panel). Homogeneity of loaded proteins (15 µg per lane) was verified by Coomasie brilliant blue staining (lower panel). **(d)** Fire blight disease incidence (± standard deviation) 21 days after *E. amylovora* infection and phenotypic assesment by AUDPC calculation over these 21 days. Each point represents the AUDPC for individual plants (52<n<70 from two independant biological repeats). Diamonds represent the mean AUDPC for each condition. For all quantified parameters, significance was evaluated with a Kruskall-Wallis test followed by a Dunn test, different letters indicating significant differences (P<0.05).

Next, to determine if *MdAGGs* are involved in the ability of greenhouse-grown apple to mount an effective ASM-induced resistance response toward *Ea*, mutant lines *mdagg-1* and *mdagg-2* were acclimatized in greenhouse, treated with ASM and further inoculated with *Ea*.

WT plants, either treated with water or ASM, were included as susceptible and resistant controls, respectively. Targeted gene expression and protein production were also analyzed in the remaining leaf tissues cut out during the inoculation procedure. *MdAGGs* expression was monitored by RT-qPCR with primers O (Table S1) designed to allow amplification of transcripts only in absence of edition at the first target site (Fig. S1c) because this site seemed to be very effective for MdAGGs edits (Fig. 1). As expected, the results show that *MdAGGs* expression was significantly upregulated by the ASM treatment compared with the water treatment in untransformed control plants. Interestingly, no significant difference in *MdAGGs* transcript accumulation was observed between the susceptible control and the ASM-treated mutant lines (Fig. 3b), suggesting that these mutant lines are unable to accumulate functional *MdAGGs* transcripts in response to ASM. Protein accumulation analysis by western blot confirmed loss-of-function for *mdagg-1* and *mdagg-2* mutants in sofar MdAGGs proteins were only detected in the resistant control (Fig. 3c). After inoculation with *Ea*, fire blight disease symptoms were scored every 3 to 4 days during 3 weeks and resistance was evaluated by determining disease incidence at the end of the experiment or disease severity by AUDPC calculation. As expected, the ASM treatment reduced by more than 50 % disease incidence and severity for the WT. In contrast, although disease incidence and severity in ASM-treated *mdagg-1* and *2* lines were both significantly lower than for the susceptible control, they were significantly higher than for the resistant control (Fig. 3d). This shows that despite ASM elicitation, *mdagg-1* and *mdagg-2* greenhouse-grown mutants are less resistant to fire blight than the resistant control.

Overall, these results demonstrate that the expression and accumulation of functional MdAGGs are crucial for apple to mount a fully effective resistance response induced by ASM.

### Molecular and phenotyping characterization of pPPO16::MdAGG10 inducible lines under *E. amylovora* inoculation challenge

Six of the nine lines obtained after *A. tumefaciens* transformation, called *pPPO16::MdAGG10-1* to *pPPO16::MdAGG10-6,* were analyzed to determine if bacterial infection with *Ea*, known to activate *pPPO16* promoter, effectively led to overexpression of *MdAGG10*. Transcript accumulation of *MdAGG10* was assessed by RT-qPCR at 24 and 40 hours post infiltration (hpi) in mock- or *Ea*-infiltrated leaves of *in vitro* shoot cultures. The untransformed ‘Gala’ variety (WT) was used as a control. Differential gene expression analysis shows that untransformed plants did not overexpress *MdAGG10* in response to *Ea* infection whereas the transformed lines did, with a log_2_ ratio (compared to the WT line) varying from 1 to 5 depending of the lines and the day of sampling (24 or 40 hpi - Fig. 4).

**Fig. 4.**
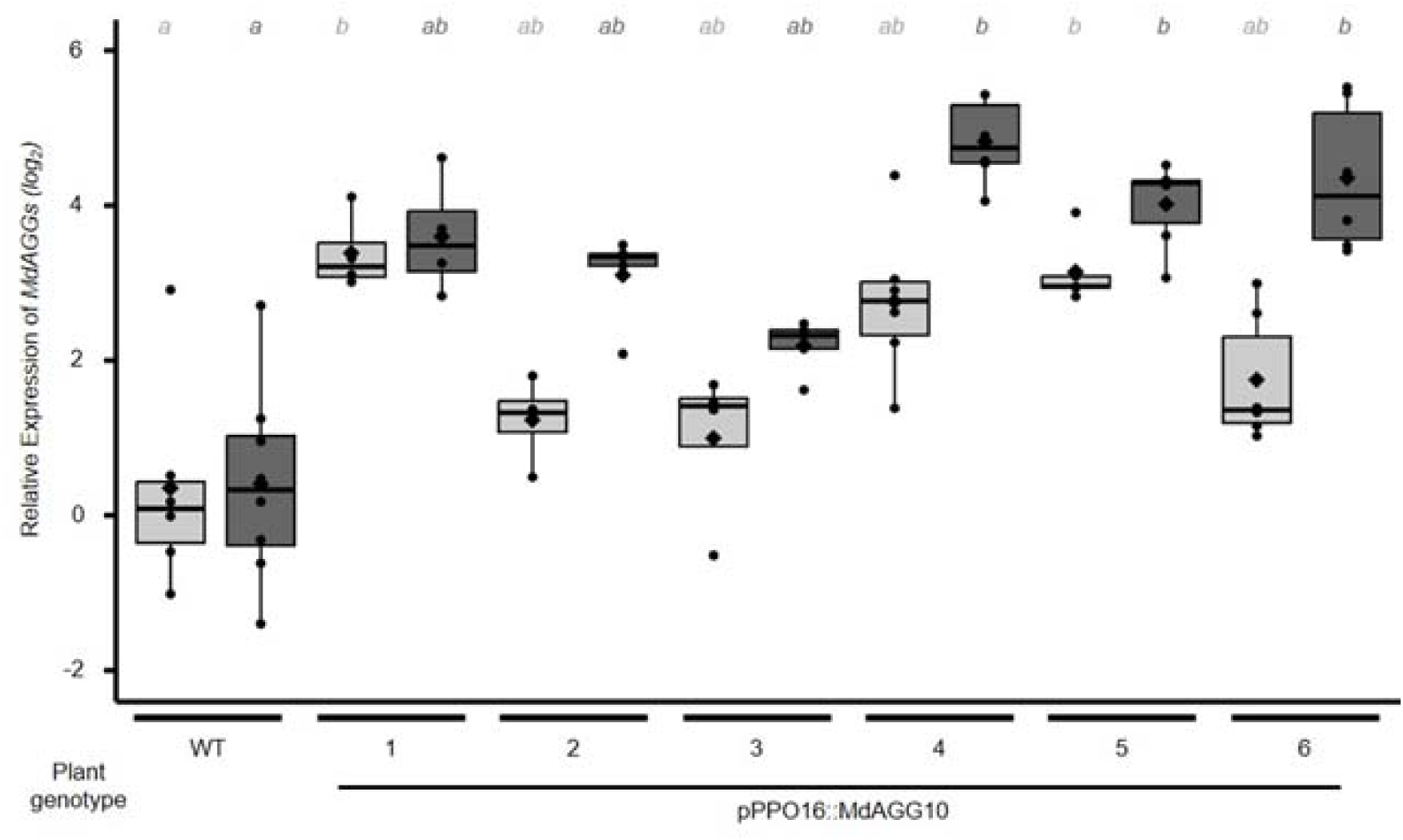
Fire blight inducible *PPO16* promoter upregulates *MdAGG*s expression upon *Erwinia amylovora* infection on *in vitro* culture shoots of *pPPO16::MdAGG10* transgenic lines. Light (respectively dark) grey boxplots represent *MdAGG10* expression 24 (respectively 40) hours after *E. amylovora* infection versus water infiltration (mock). Each point represents the *MdAGG10* expression (log_2_ ratio) in individual inoculated samples relative to the mean value of *MdAGG10* expression in the water-infiltrated biological repeats at the same day of sampling. Means, medians, interquartile range (IQR) are indicated by diamonds, lines, boxes, respectively. Each biological replicate is a pool of all leaves sampled from two independant plants (6<n<8). Significance was evaluated with a Kruskall-Wallis test followed by a Dunn test, different letters indicate different significance (P<0.05). Boxes with same letter and same colour represent medians that are not significantly different (P<0.05).

To determine if conditional upregulation of *MdAGG10* enhances apple resistance toward *Ea*, *in vitro* shoot cultures of *pPPO16::MdAGG10-1* to *6* lines were inoculated with *Ea* and disease symptoms were scored daily over 12 days following inoculation. Resistance was evaluated by determining disease incidence at the end of the experiment or disease severity by AUDPC calculation. The WT plants, either treated with water or ASM, were included as a respectively susceptible and resistant controls. Water-treated WT plants displayed significantly higher disease incidence and severity scores than ASM-treated WT plants. Fire blight incidence was significantly higher than in the susceptible control in four of the six lines (*pPPO16::MdAGG10-2* to -*4* and *5*), and intermediate to those of resistant and susceptible controls in *pPPO16::MdAGG10-1* and *6*, although more similar to incidence measured for the susceptible control (Fig. 5). In four of the six lines (*pPPO16::MdAGG10-1* to -*3* and -*6*), disease severity was significantly intermediate to those of resistant and susceptible controls, whereas in *pPPO16::MdAGG10-4* and -*5*, it was equivalent to that of the susceptible control (Fig. 5). In brief, except line *pPPO16::MdAGG10-4* and -*5*, most of *pPPO16::MdAGG10* lines *in vitro* shoots were less severely affected by fire blight than the susceptible control. Lines *pPPO16::MdAGG10-1* and *pPPO16::MdAGG10-6* were further analyzed in greenhouse.

**Fig. 5.**
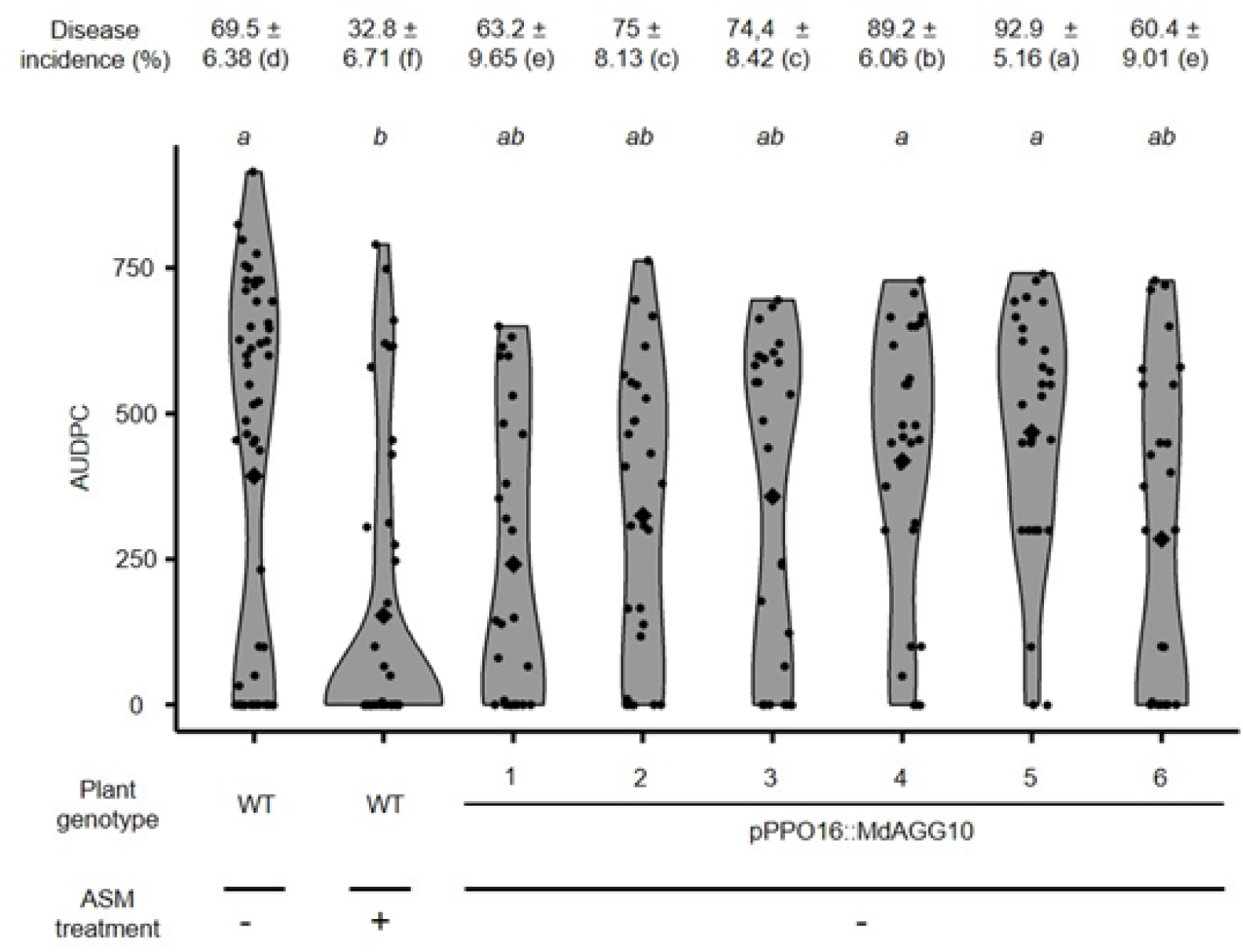
*In vitro* culture shoots of *pPPO16::MdAGG10* lines are less susceptible to *Erwinia amylovora.* Fire blight disease incidence (± standard deviation) 12 days after *E. amylovora* infection and phenotypic assesment by AUDPC calculation over these 12 days. Each dot represents the AUDPC of one shoot (29<n<35). Each diamond represents the mean AUDPC of each condition. Significance was evaluated with a Kruskall-Wallis test followed by a Dunn test, different letters indicate different statistical classes (P<0.05).

We next investigated whether infection-inducible *MdAGG10* expression provides effective resistance toward *Ea* lines *pPPO16::MdAGG10-1* and *6* grown in greenhouse and if ASM further enhances this resistance. Plants were therefore treated with ASM or water and inoculated with *Ea*, disease symptoms were scored every 3 to 4 days during 3 weeks and resistance was evaluated by determining disease incidence at the end of the experiment or disease severity by AUDPC calculation. The untransformed susceptible variety ‘Gala’, either treated with water or ASM, was included as a respectively susceptible and resistant control. Nineteen days after inoculation, plants of water-treated *pPPO16::MdAGG10-1* and *6* lines were obviously less affected by fire blight compared to the susceptible control. Disease incidence reached almost 94% in the susceptible control, which results in the stunted phenotype of plants systemically affected by *Ea* (Fig. 6a). In contrast, significantly lower disease incidence of the water-treated *pPPO16::MdAGG10-1* and *6* plants is evidenced by actively growing plants for both of these transgenic lines (Fig. 6a, b). Regarding disease severity, water-treated *pPPO16::MdAGG10-1* and *6* lines were intermediate to those of resistant and susceptible controls. Plants of both transgenic lines that were previously treated with ASM displayed lower disease incidence and severity than the resistant control (Fig. 6b). Altogether, these results show that infection-induced *MdAGG10* expression alone provides effective resistance to *Ea* in greenhouse-grown plants, with ASM treatment offering additional protection, reducing disease symptoms by approximately 90%.

**Fig. 6.**
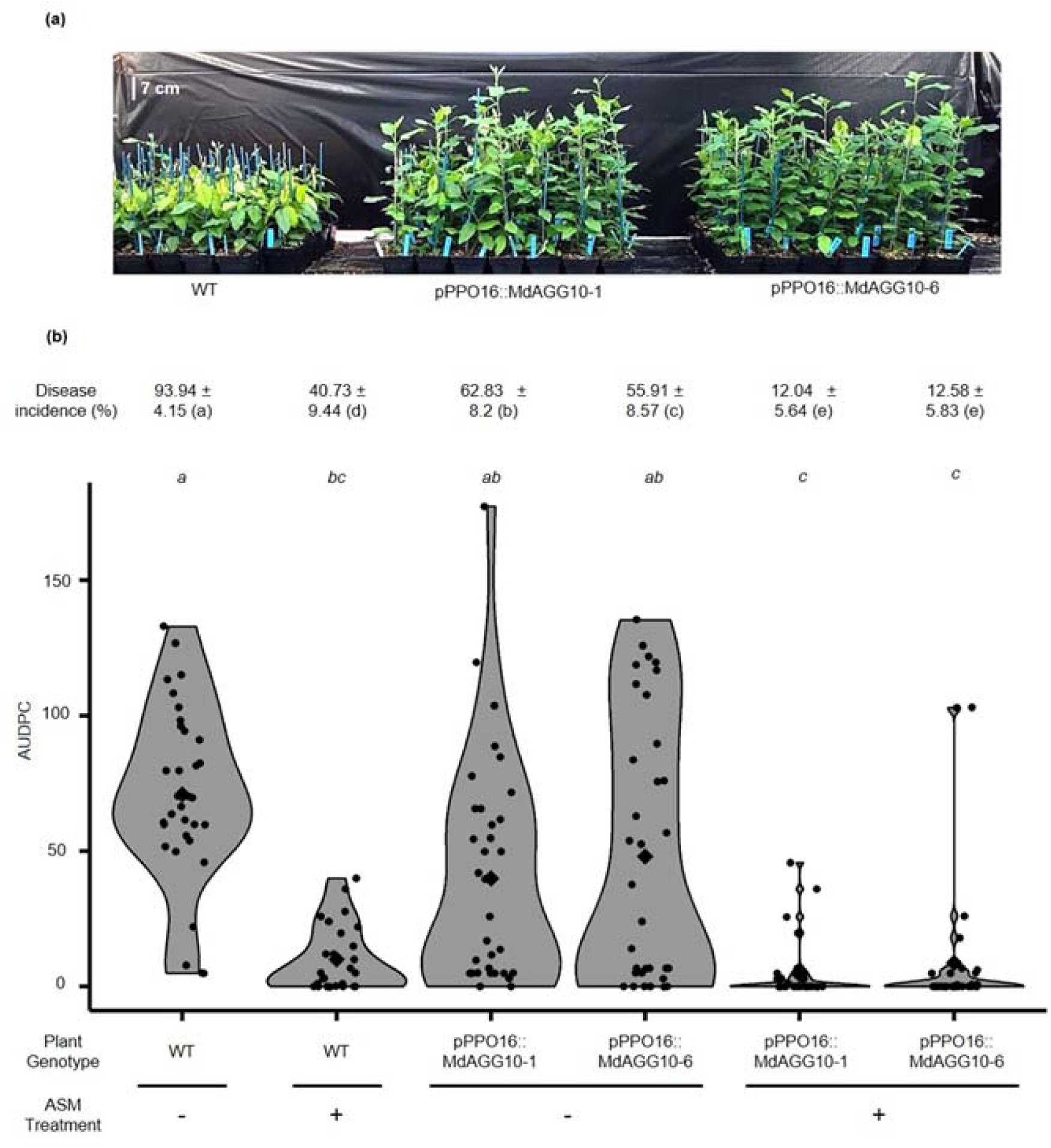
Greenhouse-grown *pPPO16::MdAGG10* apple lines are less susceptible to *Erwinia amylovora* **(a)** Representative picture of water-treated apple lines inoculated with *E. amylovora,* 19 days post-inoculation **(b)** Fire blight disease incidence (± standard deviation) 19 days after inoculation and phenotypic assessment by AUDPC calculation over these 19 days. Each dot represents the AUDPC of one plant (29<n<35). Each diamond represents the mean AUDPC of each condition. Significance was evaluated with a Kruskall-Wallis test followed by a Dunn test, different letters indicate different statistical classes (P<0.05).

## Discussion

Genetic improvement by NBT of apple varieties susceptible to fire blight have already been performed by transferring R genes from wild relatives, or knocking-out of S-genes. Here, we provide evidence that intragenic association of a disease-induced promoter with a downstream defense gene constitutes a supplementary strategy for the biotechnological toolbox aiming at enhancing and studying fire blight disease resistance in apple.

From a methodological point of view, being able to use apple shoots cultivated *in vitro* for pre-screening of transformed lines, before their acclimatization in a greenhouse, can save valuable time and money. In this work, although the observed effects were weak, they nonetheless allowed a pre-screening, and the acclimated lines in the greenhouse confirmed and reinforced the results, as already observed in others studies (Garcia-Almodovar *et al*., 2015).

Still from a methodological point of view, the two gRNAs designed for targeting all *MdAGG* members of the family, and the tandem array organization of *MdAGG1* to *MdAGG15* on chromosome 10 (Warneys *et al*., 2018) allowed large DNA deletions of several kb eliminating numerous *MdAGG* sequences in their entirety. CRISPR/Cas9 is a powerful and precise method to induce targeted mutagenesis, but it is known to create editing chimerism in plants whose transformation involves somatic embryogenesis potentially involving several cells at the origin of the regenerated line, which is the case for apple (Charrier *et al*., 2019). Rounds of adventitious regeneration step can efficiently reduce chimerism (Malabarba *et al*., 2020) but stay time-consuming. As already demonstrated in Pompili *et al*. (2020), HTS (High throughput sequencing) targeted resequencing methods can be used to capture this edition diversity in first-generation edited apple lines, both in qualitative and quantitative terms. As an initial strategy, we opted here for a partial qualitative sequencing analysis of the edits by examining a few alleles of the targeted sequences. This approach nevertheless provided a solid overview of the different types of edits involved, especially the large deletions, which would have been missed by HTS targeted resequencing due to the absence of the target sequences. In the mutant lines *mdagg-1* and -*2*, selected for further study, the gRNA1 targets consisting of *MdAGGs* target sites and the *MdDiAGG4* off-target site, were then analyzed quantitatively using an easy-to-use and effective qPCR-based method, confirming that these lines were suitable for robust greenhouse experiments.

We first demonstrated *in vitro* that most ASM-treated *mdagg* lines showed less resistance to fire blight compared to the resistant control. Mutant *mdagg-4* was the only line whose resistance was comparable to WT plants, despite editions identified within each clone analyzed. It remains unclear whether these edits affect cell types that are not directly involved in the initial defense against *Ea*, or if certain edits—such as the 212 kb deletion observed in one clone—enhance resistance by eliminating unidentified susceptibility factors. An in-depth analysis of this mutant line remains to be performed to explain its inconsistent phenotype. For the mutants selected for greenhouse assays, the decrease in resistance to *Ea* correlated with the absence of MdAGGs expression and accumulation. To the best of our knowledge, it is the first time that amaranthin-like lectins are demonstrated to be involved in plant resistance against a bacteria by loss-of-function in the native species, rather than through ectopic expression (Cabrales-Orona *et al*., 2022). Here, MdAGGs are lectins containing a unique amaranthin-like domain, able to agglutinate bacterial cells *in vitro* but devoid of proven bactericidal effect (Bodelot *et al*., 2023). Whether MdAGGs act by physically slowing bacterial progression within apple tissues thanks to their agglutination properties or by targeting other plant-derived antimicrobial compounds toward the bacterial cells remains to be investigated.

Once demonstrated that MdAGGs are necessary to mount an effective ASM-triggered resistance response to fire blight, we wanted to determine if gain-of-function by inducible-expression of MdAGG10 improves resistance to *Ea*. Because of its high expression after ASM treatment and its closest proximity to the consensus sequence of the 17 members of the family (Warneys *et al*., 2018), we choose *MdAGG10* as the best candidate to select, to promote fire blight resistance. A previous attempt performed by constitutive overexpression of *MdAGG10* did not meet our expectations in sofar overexpressing lines were as susceptible to *Ea* as wildtype ones (Bodelot *et al*., 2023). Here, our results showed that *MdAGG10* exhibited strong expression in the early stages of *Ea* infection, thanks to its association with the *Ea*-inducible promoter *pPPO16* (Gaucher *et al*., 2022), and this strong expression was associated with enhanced resistant to the disease. Moreover, combining inducible expression of *MdAGG10* and treatment with ASM led to a drastic reduction of the disease. These results, when compared to those from the previous study involving constitutive overexpression of the same *MdAGG10* gene (Bodelot *et al*., 2023), underscore the advantage of prioritizing inducible expression over constitutive expression when the aim is to enhance the activity of genes of interest. Inducible and transient expression of *MdAGG10* had here no negative impact on plant development and conferred partial resistance, whereas its constitutive CaMV p35S-driven overexpression led to pleiotropic effects on plant growth without enhancing resistance (Bodelot *et al*., 2023). Such pleiotropic effects have already been documented in studies dedicated to constitutive overexpression of lectin-encoding genes (Cabrales-Orona *et al*., 2022). The past studies which demonstrated that the ectopic expression of the amaranthin encoding gene from *Amaranthus caudatus* protects recipient plants from aphids without pleiotropic effects were performed using phloem-specific promoters (Wu *et al*., 2006). It seems therefore that amaranthin-like encoding genes can be engineered to promote plant resistance but their regulation in space and time has to be considered cautiously to avoid advert effects on plant fitness. Using native pathogen- or pest-inducible promoters like *pPPO16* to drive amaranthin-like lectin encoding genes, as we did here, could be an interesting option to test in other systems.

Until now, breeding apple resistance has focused on resistance QTLs (Emeriewen *et al*., 2021), which underlying genes turn out to be mainly R-like genes involved in gene-for-gene recognition and triggering of ETI. However, the ongoing plant-pest arms race often results in the selection of mutations in the pest’s genome. These mutations alter the structure of its avirulence proteins, allowing it to bypass the plant’s R genes and evade recognition. Bypasses of resistance QTLs to *Ea* have already been observed in apple (Emeriewen *et al*., 2018). Here, we demonstrate that it is possible to create a resistance in apple by focusing on a downstream defense gene, *MdAGG10*, which interacts with *Ea*’s LPS and EPS, possibly through electrostatic interactions (Chavonet *et al*., 2022). Indeed, the most important feature seems to be the acidic nature (*ie*. negatively charged) of the polysaccharide (Chavonet *et al*., 2022). Mutants of *Ea* engineered to produce uncharged amylovoran (the principal component of *Ea*’s EPS) were shown to be non-pathogenic on apple, phenocopying bacterial mutants that do not produce amylovoran at all (Wang *et al*., 2012). Altogether, these studies suggest that the downstream defense protein MdAGG10 targets bacterial features under strong evolutionary constraints, making the underlying apple gene a good candidate for breeding durable resistance.

## Conclusion

To conclude, in this work, we have validated MdAGGs as important compounds of apple’s defense toolbox against *Ea* and created a partial resistance to fire blight thanks to the intragenic association of the disease-inducible *pPPO16* promotor and the coding sequence of *MdAGG10*. But *pPPO16::MdAGG10* plants analyzed here do not meet the standards of intragenic plants: 1) the terminator of this construct is of viral origin and 2) the corresponding T-DNA does not have a recombination system such as in the PMF1-GFP-dest plasmid (Schaart *et al*., 2010), which allows the elimination of other elements (selection gene, recombination system itself) than the construct of interest after transformation. Functional intragenic apple lines could be obtained using the Rubisco terminator, which has already been employed in apple (Joshi *et al*., 2011). Combined with the overmentioned recombination system, such intragenic lines allowing *MdAGG10* expression upon *Ea* infection, could be upgraded toward higher resistance by pyramiding more genes involved in partial resistance. Further combined with other control methods, such as treatments with environment-friendly plant resistance inducers, will contribute to generate robust and high-level resistance enabling better control of fire blight.

## Material and Methods

### 1. Bacterial strains, culture and suspensions preparation

*A. tumefaciens* strain EHA105 containing plasmids of interest was grown in Luria Broth (LB) media (Duchefa Biochemie, DH Haarlem, Netherland) at 28°C. Suspensions of *A. tumefaciens* were prepared in liquid apple regeneration medium consisting of MS medium containing 22.7 µM thidiazuron (TDZ), 2.15 µM IBA and 100 µM acetosyringone. *Ea* strain CFBP1430 was grown on King’s B media (King *et al*., 1954) at 26°C. Suspensions of *Ea* were prepared in sterile water from solid exponential growth phase culture.

### 2. Construction of vectors and generation of transgenic lines

Binary vector CRISPR-MdAGGAll used in this study (T-DNA in Fig. S1a) derived from pDE-CAS9Kr vector (Charrier *et al*., 2019). This construct contained two gRNAs with a different promoter (gRNA1 with MdU3 and gRNA2 with MdU6) targeting the 18 MdAGG members of the family (Table 1). Each gRNA cassette marked out by attB gateway sites was synthesized independently by Integrated DNA Technology, Inc. (San Jose, CA, USA) and then cloned in pDONR207 vector by BP cloning (Gateway system; Thermo Fisher Scientific, Waltham, MA, USA). Then the ‘U6gRNA2’ cassette was placed after the ‘U3gRNA1’ cassette by restriction/ligation at XhoI/PstI sites in the donor vector and SalI/PstI sites in the destination vector, to create pDONR207-U3gRNA1-U6gRNA2 vector. Gateway LR cloning between pDONR207-U3gRNA1-U6gRNA2 vector and pDE-CAS9Kr vector was then performed to create CRISPR-MdAGGAll. Primers used to verify the cloning at each step (primers A: BP cloning and gRNAs addition, primers B: LR cloning) are indicated in Table S1. MdU3 or MdU6 promoters driving gRNAs were, respectively, found upstream MD10G1073100 and MD07G1138500 genes (https://iris.angers.inra.fr/gddh13/) by BLAST with AtU3 and AtU6 sequences (respectively, found upstream X52629 and X52528; Marshallsay *et al*., 1990). Sequences are given in Charrier *et al*. (2019). Target sequences of gRNAs were chosen with CRISPOR software (http://crispor.tefor.net/; Haeussler *et al*., 2016) using the *Malus domestica* INRAE GGDH13 Version 1.1 genome (https://iris.angers.inra.fr/gddh13/; Daccord *et al*., 2017).

Binary vector pPPO16::MdAGG10 used in this study (T-DNA in Fig. S1b) derived from pKGWFS7 (Karimi *et al*., 2002) and was constructed as followed. The promotor *pPPO16* from ‘MM106’ (MK873007, about two kilobases - kb) was amplified with Q5 High Fidelity DNA Polymerase (New England Biolabs, Ipswich, MA, USA) according to the manufacturer instructions and with primers C adding an *EcoRI* restriction site in 3’. Amplified fragment was then cloned into p-ENTR/D TOPO (Invitrogen, Carlsbad, CA, USA) and then transferred into the destination vector pKGWFS7 by Gateway recombination to create pKGWFS7-pPPO16. *MdAGG10* CDS (MD10G1027210, 498 base pairs) was also amplified with the Q5 high fidelity DNA polymerase according to the manufacturer instructions and with primers F adding *EcoRI* and *NcoI* restriction sites in 5’ and 3’ respectively, then cloned into pGEM-T easy (Promega, Madison, WI, USA) according to the manufacturer instructions. The *EcoRI*-MdAGG10-*NcoI* fragment was then inserted in pKGWFS7-pPPO16 by restriction/ligation to create pPPO16::MdAGG10. Primers used to clone or to verify the cloning at each step (primers C to H) are indicated in Table S1.

Apple transgenic lines were generated as previously described in Malabarba *et al*. (2020).

### 3. Plant growth conditions and ASM treatment

*In vitro* apple shoots were micropropagated as in Malabarba *et al*. (2020) and the rooting conditions were the same as those reported in Faize *et al*. (2003). The rooted shoots were transferred into greenhouse and grown under 22°C, humidity rate of 80% and shading of 500 W.m^-^² during 8 weeks after acclimatation.

*In vitro* shoots and greenhouse grown plants are treated with a 0.2 g.L^-1^ ASM (Bion® 50WG, Syngenta, Basel, Switzerland) solution 3 days before inoculation, with either an Ecospray sprayer (Seidden, Madrid, Spain) for *in vitro* shoots or a 750 mL hand sprayer for greenhouse grown plants. The ASM solution was filter-sterilized at 0.22 μm (Pall Corporation, Port Washington, NY, USA) before spraying the *in vitro* shoots.

### 4. Plant inoculation, phenotyping and sampling

Apple *in vitro* shoots and greenhouse grown plant inoculations were performed with a 10^7^ colony forming unit.mL^-1^ *Ea* suspension prepared in sterile water. Both *in vitro* shoots and greenhouse grown plant were inoculated with scissors previously dipped in *Ea* suspension, by cutting the lower third of one of the apical leaf for *in vitro* shoots, or of the last fully developed leaf for greenhouse plant. Cut parts of leaves from two different plants were then pooled and frozen at -80°C for subsequent analysis. Fire blight incidence was measured by checking the presence of necrotized cells beyond the inoculated leaf petiole. Fire blight severity corresponded to the Area Under Disease Progression Curve (AUDPC - Shaner and Finney, 1977) calculated with fire blight symptoms measured differently on *in vitro* shoots or greenhouse grown plants. On greenhouse grown plants, they were measured 4, 7, 11, 14 and 19 day post inoculation (dpi) by measuring the length of necrotized shoot whereas on *in vitro* shoots, a score has been assigned at 4, 5, 6, 7, 8 and 10 dpi according to the proportion of necrotized tissues observed on each plant. These scores are summarized in Table S2.

### 5. Nucleic acids extraction and reverse transcription

Genomic DNA (gDNA) was extracted as described in Fulton *et al*. (1995). RNA extraction, reverse transcription and MdAGGs expression analysis were performed as described in Bodelot *et al*. (2023).

### 6. Genotyping of transgenic lines by PCR and targeted DNA sequencing

T-DNA insertion verifications with primers B, H, I, J, K, L (Table S1), were performed by PCR amplifications with GoTaq G2 Flexi DNA polymerase (Promega) according to manufacturer’s recommendations. The PCR reaction conditions were: 95 °C for 5 minutes followed by 35 cycles at 95 °C during 30 seconds, 58 °C during 45 seconds and 72 °C during 60 seconds with a final extension of 5 minutes at 72 °C. PCR products were separated on a 2% agarose gel.

For the qualitative characterization of MdAGGs edits, we used primers M (Table S1) flanking the sequence containing the target zones of the two gRNAs. Amplifications were performed with Q5® High Fidelity 2X MasterMix according to manufacturer’s recommendations. The PCR reaction was performed with an initial step at 98°C during 30 seconds followed by 40 cycles at 98°C during 10 seconds, the appropriate melting temperature during 30 seconds and 72°C during 60 seconds with a final extension at 72°C during 120 seconds. Amplicons were cloned with a CloneJET PCR kit (Thermo Fisher) according to manufacturer’s recommendations, and bacterial transformation was performed with Top10 competent *E.coli* cells subsequently spread on LB medium complemented with 100 mg.mL^-1^ ampicilin for 24 hours at 37°C. Bacterial clones containing MdAGG amplicons were selected by on-colony PCR using N primers (Table S1) present on the plasmid and framing the insertion zone, with GoTaq G2 Flexi DNA polymerase according to manufacturer’s recommendations. Selection was on the basis of the expected size of the amplicons inserted in the event of slight modifications of the gRNAs targeted sequences (indels or substitutions) or of elimination of the portion of sequence between the two gRNAs targeted sequences. For each transgenic line, 4 bacterial clones containing putative edited MdAGGs sequences were sent for sequencing with the reverse N primer (Table S1) to Azenta Life Science company (Burlington, MA, USA). The resulting sequences were aligned with the CDS of *MdAGG1* to *MdAGG18* from the apple GDDH13 genome sequence (Daccord *et al*., 2017) using the MultAlin software (Corpet, 1988).

### 7. qPCR analysis

*MdAGG* expressions and quantifications of *MdAGGs* and *MdDiAGG4* edition rates in *mdagg-1* and *mdagg-2* lines were measured by qPCR. Briefly, cDNAs (expressions) or genomic DNA (edition rates) were mixed with MESA Blue 2X PCR MasterMix for SYBR Green Assays with fluorescein (Eurogentec, Liege, Belgium) and primers (O for expressions, P and Q for edition rates; Table S1) at the appropriate concentrations. Reactions were performed on a CFX Connect Real-Time System (Bio-Rad Laboratories, Hercules, CA, USA) for the qPCR and melt curves were done to check the absence of non-specific amplifications and primer-dimers products. Relative expressions and edition rates quantifications were calculated using the 2-ΔΔCt method. The normalization factor was calculated with 3 housekeeping genes (Actin, GAPDH and TuA; primers R, S, T respectively in Table S1) as recommended by Vandesompele *et al*. (2002).

### 8. Protein extraction, separation and immunodetection

Proteins were extracted from frozen leaves as described in Chavonet *et al*. (2022). Protein separation and immunodetection were done as described in Bodelot *et al*. (2023).

### 9. Data analysis

Data analysis was performed using Rstudio software (R Core Team, 2023) and the graphical representations were generated using the packages “ggplot2” (Wickam, 2016) in association with “ggpubr” (Kassambara, 2023). Fire blight incidence on apple shoots was generated with the package “boot” (Canty and Ripley, 2024). A non-parametric test (Kruskall-Wallis) followed by a post-hoc Dunn test were used for all the experiments, using packages “FSA” (Ogle *et al*., 2023) in association with “rcompanion” (Mangiafico, 2024).

## Supporting information

Supplemental Informations

## Acknowledgment

The authors are grateful for the technical support provided by the ANAN shared platform of the SFR QUASAV and the PHENOTIC platform (greenhouse facilities). The authors thank their funders: these researches were funded by the Agg-FREDI project (INRAE BAP Department 2020-2021) and a PhD grant was awarded to AB by INRAE-BAP Department and the Pays-de-la-Loire Region.

## Author contributions

The design of the research was performed by EV and AD. Performance of the research: apple transgenic lines were obtained by AB, ND, ER and EV. Plant inoculation and phenotyping were performed by AB, ND, ER, CH, EV and AD. Genotyping, qPCR analysis, immunodetection were completed by AB, AD and EV. Data analyses were completed by AB, AD and EV. Interpretation and writing of the manuscript: the manuscript was drafted by AB, completed by EV, AD and MNB and reviewed by all the authors. EV, AD and MNB supervised the research project.

## Data availability

All data supporting the findings of this study are available in the article and in its supplemental information files.

## Competing interests

The authors declare no competing interests.

## Supplementary Material

Supplementary Material may be found online.

**Fig. S1** T-DNA constructs and positions of gRNAs targets, quantitative PCR primers, and Western-blot probes, on MdAGG sequences.

**Fig. S2** Genotyping of apple transgenic lines.

**Table S1** Primer used in this work.

**Table S2** Disease score assigned for AUDPC calculation on in vitro culture shoots inoculated by Erwinia amylovora.

## Notes

### Competing Interest Statement

The authors have declared no competing interest.

### Summary of Updates

We have inserted the figures in the text and corrected an omission in the materials and methods.

