## Supplemental Informations for "MdAGG apple lectins in Fire Blight resistance: CRISPR/Cas9 validation and their potential for intragenesis approaches"

The following Supporting Information is available for this article:

**Fig. S1** T-DNA constructs and positions of gRNAs targets, quantitative PCR primers, and Western-blot probes, on *MdAGG* sequences.

**Fig. S2** Genotyping of apple transgenic lines. (a) PCR on genomic DNA of the five transgenic apple lines with CRISPR/Cas9 targeting *MdAGG* sequences.

**Table S1** Primer used in this work.

**Table S2** Disease score assigned for AUDPC calculation on *in vitro* culture shoots inoculated by *Erwinia amylovora*.

**Fig. S1** T-DNA constructs and positions of gRNAs targets, quantitative PCR primers, and Western-blot probes, on *MdAGG* sequences. (a) The *Cas9* gene from *Streptococcus pyogenes* is driven by *PcUbi4-2* promoter (P) from parsley (*Petroselinum crispum*) and transcription is terminated by the *Pea3a* terminator (T) from pea (*Pisum sativum*). gRNA1 and 2 are respectively driven by *MdU3* and *MdU6* promoters from *Malus domestica* and transcription is terminated by a polyT terminator (adapted from Charrier et al, 2019). (b) The *MdAGG10* gene expression from *Malus domestica* is driven by the *PPO16* promotor and transcription is terminated by the *CaMV35S* terminator. (a) & (b) Transformants are selected with a *nptII* gene controlled by *nos* promoter and terminator from *Agrobacterium tumefaciens*. *AttB1* and *2*: sites resulting from the Gateway® LR recombination. LB and RB: T-DNA borders. (c) Schematic representation, on a *MdAGG* sequence - in blue with targets of the gRNAs in green - of the position of primer pairs used in quantitative PCR to quantify the editing rate at RNA level in CRISPR lines (in yellow, pair P in Table S1) and to quantify the expression level of *MdAGG*s in CRISPR or inducible lines (in white, pair O in Table S1). (d) Schematic representation, on a MdAGG sequence of the positions of probes used to synthetize anti-MdAGG antibodies for western blot analyses.

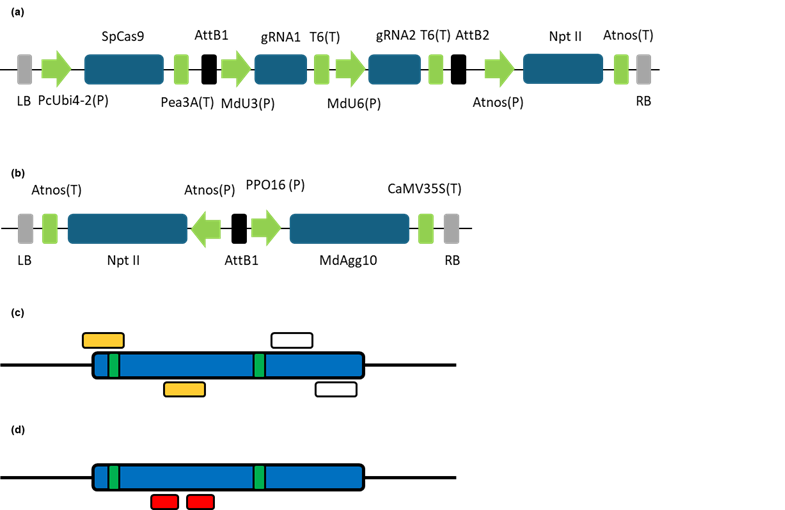

**Fig. S2** Genotyping of apple transgenic lines. (a) PCR on genomic DNA of the five transgenic apple lines with CRISPR/Cas9 targeting *MdAGG* sequences. Lane 1 to 5: transgenic *mdagg* lines; (b) PCR on genomic DNA of the nine transgenic apple lines with *pPPO16::MdAGG10* construct. Lane 1 to 9: pPPO16::MdAGG10 transgenic lines. AgB: Plasmid DNA extracted from the *A. tumefaciens* strain used for transformation as a positive control of T-DNA presence and a negative control of lines Agrobacterium contamination; W: water template as a PCR control ; WT : total DNA extracted from the ‘Gala’ non transformed genotype, as a negative control of T-DNA presence.

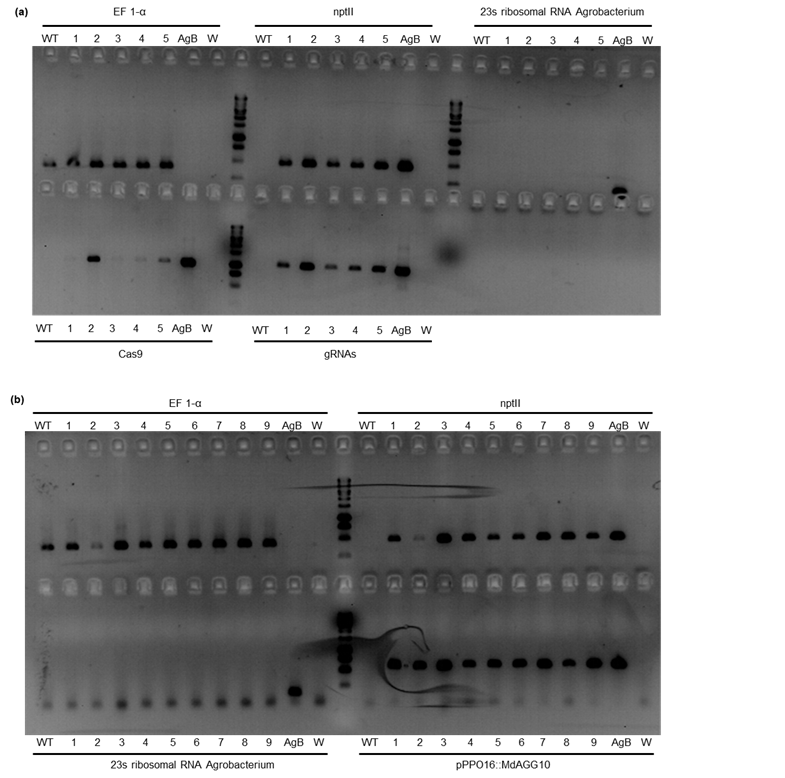

**Table S1** Primer used in this work. * « MD » accessions are available at https://iris.angers.inra.fr/gddh13, within the "curated CDS" track.

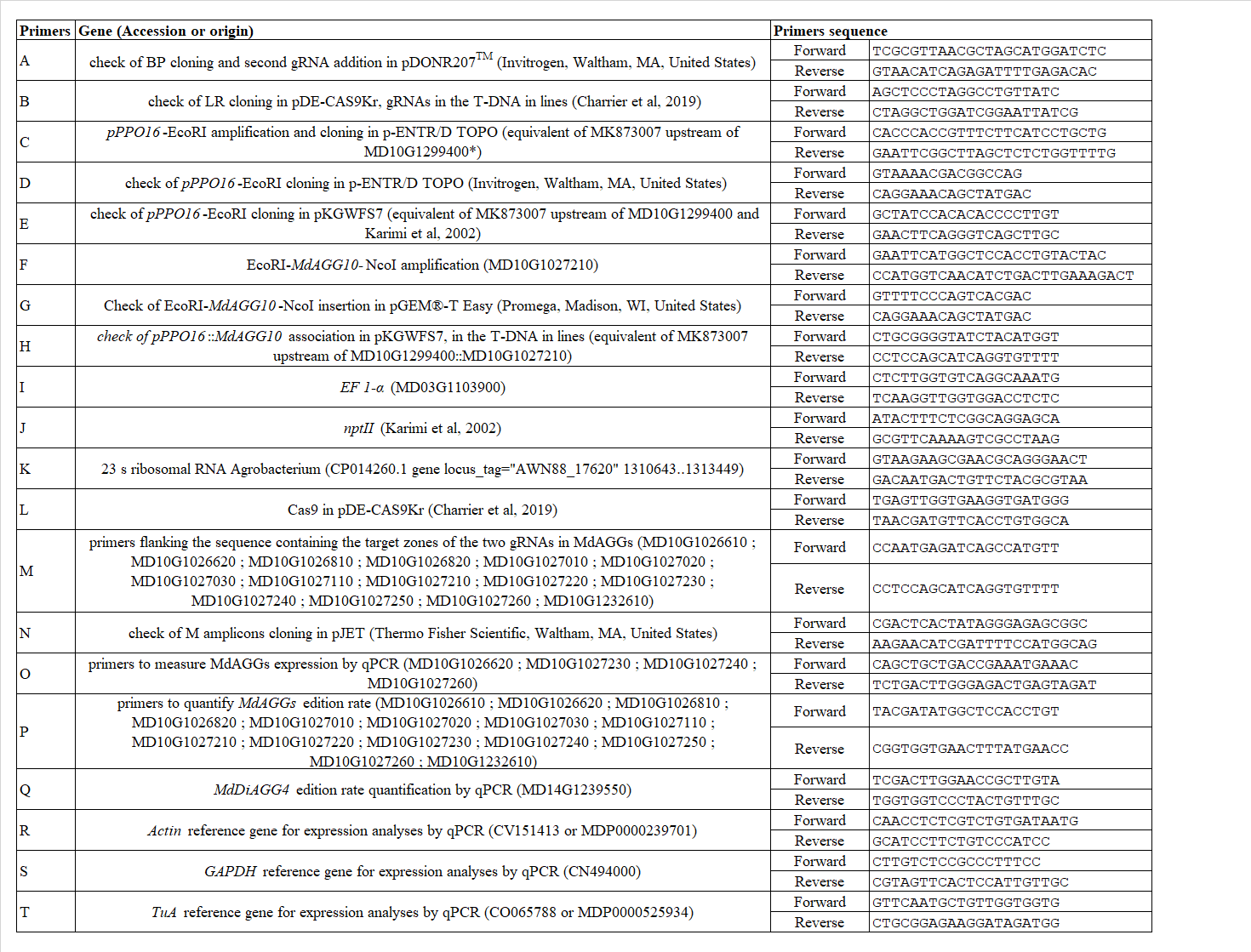

**Table S2** Disease score assigned for AUDPC calculation on *in vitro* culture shoots inoculated by *Erwinia amylovora*.

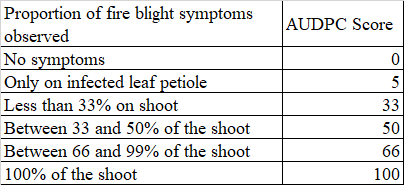
